# Advancing marker-gene-based methods for prokaryote-mediated multifunctional redundancy: exploring random and nonrandom extinctions in a watershed

**DOI:** 10.1101/2024.03.14.584931

**Authors:** Wan-Hsuan Cheng, Takeshi Miki, Motohiro Ido, Kinuyo Yoneya, Kazuaki Matsui, Taichi Yokokawa, Hiroki Yamanaka, Shin-ichi Nakano

## Abstract

Multifunctional redundancy, the extent of loss in multiple ecosystem functions with decreasing biodiversity, stands as a crucial index for evaluating ecosystem resilience to environmental changes. We aimed to refine a marker-gene-based methodology for quantifying multifunctional redundancy in prokaryotic communities. Using PICRUSt2, we predicted KEGG orthologs (KOs) for each Amplicon Sequence Variant (ASV), assessed community-wide KO richness, and validated predictions against experimentally quantified phenotypic multifunctionality. Additionally, we introduced a refined regression on ASV richness–KO richness curves, providing a reliable estimate of the power-law exponent within computational time constraints, serving as the multifunctional redundancy index. Incorporating various non-random extinction scenarios alongside a random one allowed us to quantify estimate variations between scenarios, providing conservative estimates of multifunctional redundancy. Applied to Lake Biwa and four of its inlet rivers, the refined methodology unveiled spatio-temporal variations in multifunctional redundancy. Our analysis demonstrated lower redundancy in Lake Biwa compared to rivers, aiding in prioritizing conservation targets and inferring distinct community assembly processes. Future directions include a deeper exploration of KO composition information for detailed multifunctionality quantification and the refinement of extinction scenarios. This study demonstrates the promising integration of bioinformatic functional prediction and modeling biodiversity loss, offering a valuable tool for effective ecosystem management.

## Introduction

Microbial diversity is pivotal for sustaining ecosystem functions across both aquatic and terrestrial environments, spanning from local to global scales (1,2). However, this critical role faces challenges posed by the risks of local extinction of microbial species (3,4) and spatial homogenization of microbial community composition (5,6).

In assessing the resilience of ecosystems to environmental changes and disturbances, the extent of biodiversity loss and its subsequent impact on ecosystem functions are crucial factors (7–10). Functional redundancy, representing the degree of overlap in ecosystem functions among taxonomic units within a community, is a critical concept in addressing the latter concern. Ecosystems characterized by lower functional redundancy are more vulnerable to disturbances, highlighting the necessity for prioritized conservation efforts.

Functional redundancy is originally defined as the degree of maintaining a single ecosystem function in the face of taxonomic richness loss (11–14). Various methods for quantifying functional redundancy have emerged, largely rooted in the multifunctionality concept (9,15), which addresses the simultaneous assessment of multiple ecosystem functions. In addition to simple indices utilizing taxonomic and functional diversity measures, as comprehensively reviewed by Galland *et al.* 2020, the relationship between taxonomic richness loss and functional richness loss is depicted through simulations of random or non-random taxonomic extinctions (17,18). The shape parameter of the resulting curve, such as the area under the curve, is subsequently employed as an index of functional redundancy (18,19). While some studies focus on the correlation between taxonomic composition dissimilarity and functional dissimilarity to infer the presence and strength of functional redundancy (20), it should be noted that this discussion primarily revolves around the linkage between taxonomic composition and functional composition. This approach is not explicitly designed to predict the loss of multiple functions due to taxonomic richness loss.

Extending the shape parameter approach to microbial communities, the previous study (21) incorporated the entirety of the functional gene pools within a community as a proxy for genome-based multifunctionality. It introduced the exponent coefficient from power-law fitting on the curve of taxonomic richness and functional gene richness as an index of genomic multifunctional redundancy. In the study, Miki, Yokokawa and Matsui (2014) employed in-silico simulations with habitat-specific pseudo-communities sourced from the Microbial Genomic Database (MBGD, (22)). The study revealed that functional redundancy was considerably lower than indicated by earlier empirical studies (23). This low functional redundancy has since received support from multiple studies using different approaches (2,24,25). However, it is worth noting that quantitative comparison of multifunctional redundancy across various methods poses challenges (26), underscoring potential weaknesses in asserting unanimous support for the observed lowness in functional redundancy.

Nonetheless, it is crucial to highlight the limitations of the methodology introduced by (21), which we aim to address and improve upon in this study. First of all, this earlier work presents some issues because it concentrated on creating pseudo-communities from the Microbial Genomic Database (MBGD(22)) with a small number of naturally sampled communities. Functional overlap may be underestimated as a result of the deliberate selection of just one strain from each genus to build the pseudo-communities in order to avoid introducing phylogenetic bias. This is particularly significant when compared to natural communities that permit the cooccurrence of closely related taxonomic units, potentially biasing the estimation towards lower functional redundancy. Moreover, when applying the proposed method to marker-gene compositions from natural samples, bioinformatic tool such as PICRUSt or tax4fun (27,28) was not used to connect marker-gene information to whole genome prediction (21). Instead, the phylogenetically closest strain with whole genome information registered in the MBGD database was manually selected, which could introduce biases. Another limitation arises from the consideration of only random extinction, which might underestimate the impact compared to realistic non-random sequences of extinction. Lastly, while Ruhl *et al.* 2022 employed a similar power-law fitting approach with direct measurements of functional genes through both amplicon sequences and metagenomics, it revealed limitations in fitting the power-law curve. This cautions against generalizing findings based on power-law fitting in the context of assessing functional redundancy.

Due to the persisting cost-ineffectiveness of metagenomic sequencings and the complexity of taxonomic assignment algorism to metagenomes (30), methods that rely only on marker-gene (16S rRNA gene) data are still important for assessing multifunctional redundancy. In light of this, our study aimed to improve the methodology originally proposed by Miki, Yokokawa and Matsui (2014). Subsequently, we applied this refined, and cost-efficient method to analyze natural samples, with the objective of quantifying variations in functional redundancy across different locations within a single watershed, encompassing both spatial and temporal scales.

In our study, we focused on two key aspects to enhance the existing methodology. Firstly, for the improvement of functional prediction from marker-gene composition, we utilized PICRUSt2 (31). This approach was indirectly validated through a comparison with metabolic profiling, specifically conducted via the Ecoplate incubation experiment (32). Secondly, our study incorporated both random and non-random extinction scenarios to comprehensively estimate the range of functional redundancy. This consideration allowed us to evaluate the impact of varying extinction patterns on the estimation of functional redundancy. Furthermore, we critically assessed the fitting of power-law regression and improved the regression method to ensure robustness and reliability in our analyses.

We then applied the refined methodology to various sites within a single watershed, including Lake Biwa and its four inlet rivers in Japan. Multiple samplings were conducted at each site, enabling us to investigate variations in functional redundancy across both spatial and temporal dimensions. Additionally, by pooling all samples and simulating species extinctions as a single metacommunity (33–36), we aimed to elucidate potential mechanisms contributing to the variations in multifunctional redundancy among these five sites in the single watershed. These steps are pivotal in contributing to the ongoing debate about redundancy levels, as discussed in earlier studies (2,14,37–39). Moreover, it plays a crucial role in outlining a strategy for assessing biodiversity impact (40) and determining conservation priorities based on functional redundancy (19).

## Materials and methods

### Sample collection

We collected water samples from the surface at the Ie-1 station in the north basin of Lake Biwa (35°12’58"N, 135°59’55"E) on July 3, July 30, September 10, and October 17 2019, and four of the inlet rivers of Lake Biwa: Yasu River (35°02’35.5"N, 136°01’10.0"E), Hino River (35°06’09.1"N, 136°04’37.4"E), Echi River (35°11’44.0"N 136°10’46.5"E), and Ane River (35°24’45.1"N, 136°16’59.0"E) on July 9, August 7, September 17, and October 15 2019, respectively. We filtered 250 mL (for the river samples on July 9) or 500 mL (for all the other samples) with ϕ 0.22 μm Sterivex^TM^ cartridges (SVGV010RS, Merck Millipore Darmstadt, Germany) filled with 1 g of zirconia beads implemented (ϕ 0.5 mm, YTZ-0.5; AsOne, see Ushio 2019) for the amplicon sequencing. The variations in filtered water volumes were contingent on water sample conditions, primarily influenced by fine particles that tended to clog the filter pores. Following filtration, each filter cartridge received the addition of 1 mL of RNAlater solution. Additionally, 50 mL of unfiltered water was collected specifically for bacterial direct count and the ecosystem functioning experiment. The samples designated for both amplicon sequencing and ecosystem functioning experiments were promptly transported back to the laboratory. Throughout the transportation period (up to 6 hours), the samples were maintained at 4°C. The filter cartridges were subsequently preserved in a freezer at –20°C until DNA extraction and further processing, while the unfiltered samples were immediately utilized for ecosystem functioning experiments.

### Bacterial direct count

Bacteria were enumerated directly under an epifluorescence microscope using the SYBR Green I staining method by Honjo *et al.* (2007) with some modifications. Briefly, 1 mL of the water sample was mixed with 10 µl of 200-fold diluted SYBR Green I solution for bacterial staining. After staining for 10 minutes, the bacteria in the water sample were trapped onto a 0.2 µm pore-size polycarbonate membrane filter (Advantec, Japan) and then mounted on glass slides with a drop of immersion oil (Olympus, Japan). We randomly selected ten fields per filter, and in total, more than 300 bacteria were counted using an Olympus BX51 epifluorescence microscope equipped with an oil-immersion objective (UPlanFL 100×/1.30) lens at 100x magnification under blue excitation.

### Ecosystem functioning experiment

In this study, we measured the capabilities of processing 31 organic carbon substrates as the index of multifunctionality using the Ecoplate (Biolog, Hayward, CA, USA) (21,32). The Ecoplate is a phenotypic microarray containing triplicate wells for each single carbon substrate and three control wells with no substrate. Each well also contained tetrazolium violet dye, which turned purple when the substrate within the well was catabolized. To conduct this experiment, we inoculated 100 μl of sample water into each well of an Ecoplate and conducted the incubation of all the Ecoplates in a single cooling incubator with 20°C (As One). We incubated the Ecoplates for 14 days to ensure that the color development reached saturation. Color development was measured with an optical density (OD) microplate reader (iMark, Bio-rad) set at 595 nm every day from the 0th day to the 14th day.

To evaluate ecosystem functioning, the cumulative color development from 1^st^ day to the 14^th^ for each well of the Ecoplate was calculated for assessing the integration of the color density development curve. The resulting integrated value was normalized by dividing it by the integration period (32). Furthermore, to standardize the background turbidity and color development in relation to in situ dissolved organic carbon (DOC), the integrated value of the control well was subtracted from each integrated value of the Ecoplate. After averaging the values among triplicate wells to minimize experimental error, values for 31 different functions were obtained, which were then be used as our EF indices.

To estimate the multifunctionality based on 31 EFs derived from the Ecoplate as a proxy of phenotypic ecosystem multifunctionality (MF_P_), threshold method (Zavaleta et al. 2010) was applied as

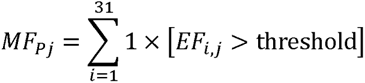

where *EF_i,j_* represents the value for ecosystem function *i* in a given community of the j^th^ observation. The threshold value corresponds to the 5% of the maximum value among 20 samples (including 7 samples that were excluded for functional gene prediction processes) for each function (Zavaleta et al. 2010).

### DNA extraction, PCR amplification and sequencing

We employed the DNeasy Blood & Tissue Kit (QIAGEN, Hilden, Germany) for DNA extraction from filter cartridges, following the protocols outlined by Miya *et al.* (2016) and Ushio (2019). Subsequently, the extracted DNA served as a template for the polymerase chain reaction (PCR) targeting the V4 region (∼250 bp) of the 16S rRNA gene, utilizing prokaryotic universal primers (515F by Parada, Needham and Fuhrman 2016 and 806R by (45). To ensure enhanced reproducibility and consistent outcomes, we implemented a two-step PCR approach (46). During both the extraction and PCR processes, we included an extraction negative control and a PCR negative control to monitor potential contamination. Sequencing of the PCR amplicons was conducted on the Illumina Miseq platform, generating 2×300 bp paired-end reads. Additional details on the experimental procedure can be found in the Supplementary Data (Supplementary Methods). The raw sequence data have been deposited in the NCBI Sequence Read Archive under the accession number PRJNA1080231.

### Sequence data processing

All pipelines used for processing sequence reads and generating amplicon compositions followed Ushio 2019. In brief, the raw MiSeq data were converted into FASTQ files using the bcl2fastq program provided by Illumina (bcl2fastq v2.18) without demultiplexing, and the FASTQ files were subsequently demultiplexed using Claident v0.2.2018.05.29 (http://www.claident.org, (Tanabe and Toju 2013)). Only reads that matched both the Illumina tag and primers were utilized for subsequent bioinformatic processes. The demultiplexed FASTQ files were analyzed using the Amplicon Sequence Variant (ASV) method implemented in DADA2 (v1.26.0) (47). Initially, primer removal was carried out using of the external software cutadapt v2.6 (Martin 2011). Subsequently, sequence quality filtering was performed with the DADA2::filterAndTrim() function, and error rates were determined using the DADA2::learnErrors() function, with the MAX_CONSIST option set to 20. Although the DADA2 algorithm typically processes each sample independently, this default approach tends to remove singletons and doubletons within individual samples, thereby impeding the estimation and standardization of sampling coverage. To address this limitation, we deviated from the default setting and combined all samples for sample inference using dada (…, pool = TRUE)(49). This modification allows the preservation of ASVs that appear once or twice in each sample (i.e., local singletons or doubletons) while eliminating ASVs that appear once or twice only in the pooled samples (i.e., global singletons or doubletons). After the removal of spurious sequences by the DADA2 algorithm, paired-end reads (i.e., overlapping by at least 20 bases) were merged into ASVs. Chimeric sequences were eliminated using the DADA2::removeBimeraDenovo() function. Taxonomic assignments of ASVs were performed using the SILVA database (version 138.1) (Quast et al. 2013). Any ASVs classified as Mitochondria or Chloroplast were subsequently eliminated.

The presence of ASVs in the negative controls may suggest potential contamination during the experiments (Supplemental Data). Therefore, the maximum abundance for each ASV observed in these negative controls was calculated and excluded from other samples. Note that if the resulting abundance was greater than zero, we assigned the value as normalized ASV abundance. In cases where the resulting abundance was not greater than zero, we set the normalized ASV abundance to zero.

To standardize the sampling coverage of samples on each sampling date, the coverage-based approach (Chao et al. 2014) was employed. Samples with coverage lower than 90% were excluded from further processing (Table S1). Among the remaining 13 samples, the minimum sample coverage (i.e., 93.32%) was calculated, and this fixed coverage was applied to subsample the ASV abundance 100 times for each sampling date. Subsequently, the average of the 100 subsampled ASV tables was utilized to depict the prokaryotic composition of microbial communities. When estimating the abundance of each ASV (cells/mL), we used the frequency distribution of ASVs (0-1) with the total bacterial count (cells/mL).

### Functional gene prediction

To predict functional genes, the representative sequence of ASV, along with a BIOM table that excluded ASVs classified as Mitochondria and Chloroplast, was employed in PICRUSt2 (31). The default PICRUSt2 pipeline utilized the Integrated Microbial Genomes (IMG) databased as week as KEGG database to generate KEGG Ortholog (KO) predictions for each input ASV. Each KO entry denotes an ortholog group associated with a gene product in the KEGG pathway diagram (50). Following the PICRUSt2 guidelines, ASVs with a nearest-sequenced taxon index (NSTI) score above 2 are typically considered as poor alignments with existing reference sequence in the IMG. Thus, these ASVs were removed from both KO compositional information and the averaged-rarefied ASV composition.

To estimate genomic multifunctionality (MF_G_) of a community, we multiplied the composition vector representing the presence/absence of ASVs, derived from the averaged-rarefied ASV composition, with the matrix indicating the presence/absence of KOs. This multiplication, performed as an inner product, resulted in a community-wise KO list in vector form. Subsequently, we calculated the KO richness for each community, serving as a proxy for MF_G_.

### ASV-extinction simulations

#### Setting for ASV-extinction simulations and fitting of power-law curves

We simulated the ASV richness loss as a proxy of taxonomic diversity (TR) loss and its subsequent impact on the KO richness loss as a proxy of genomic multifunctionality (MF_G_) loss within each local community. Our first aim was to improve the power-law regression method of this relationship (*MF_G_* = *cTR^a^*) for quantifying microbial community multifunctional redundancy. The exponent *a* indicates the degree of *MF_G_* changes with *TR*, while the coefficient *c* represents the expected *MF_G_* with a single taxonomic unit (*TR* = 1). To achieve this, we first adopted the random extinction scenario, assuming a random order of ASV extinctions within a community. This scenario served as a baseline for evaluating multifunctional redundancy and also facilitated the assessment and refinement of the power-law regression method. Rather than numerically simulating sequences of randomly ordered ASV extinctions, we employed an analytical formula for the *TR*-*MF_G_* relationship (21). Originally designed for a species accumulation curve and its rarefaction (51), this formula allows rapid quantification of the expected KO richness across a sequence, ranging from a single ASV to the maximum ASV richness (e.g., 1, 2, …, 1000), resulting in a high-resolution *TR-MF_G_* curve. Additionally, we generated a low-resolution *TR-MF_G_* curve with an arithmetic progression of ASV richness at a consistent interval (e.g., every 5% of the maximum ASV richness). It’s important to note that a low-resolution *TR-MF_G_* curve is exclusively suitable for non-random extinction scenarios, as the analytical formula is not applicable to them, and it is impractical to simulate KO richness for the entire sequence from a single ASV to the maximum ASV richness. The comparison between high- and low-resolution *TR-MF_G_* curves was expected to guide the development of a more robust methodology for evaluating multifunctional redundancy.

Using linear regression on log_10_-transformed data (log_10_(*MF_G_*) = log_10_(*c*) + *a*log_10_(*TR*)), the regression line tends to overestimate *MF_G_* levels when *TR* levels are small. This tendency is also indicated in Fig. S3 of Ruhl *et al.* 2022, where log-transformed data, being less dense with smaller TR levels, leads to overestimation. As our focus is on the initial phases of microbial extinctions and their impacts on ecosystem multifunctionality, the bias introduced by severe extinctions (e.g., 95% reduction of the maximum ASV richness) does not significantly compromise the efficacy of the power-law regression method for assessing multifunctional redundancy. However, developing an appropriate fitting procedure is also crucial to ensure quantitative comparability between high- and low-resolution *TR-MF_G_* curves, corresponding to analytical and numerical methods. To determine the specific ranges where power-law fitting on the low-resolution curve aligns reasonably with the high-resolution curve, we conducted a comparative analysis of slope estimates across three intervals: 1) from a single ASV to the maximum ASV richness, 2) from 10% to 100% of the maximum ASV richness, and 3) from 50% to 100% of the maximum ASV richness.

#### Non-random extinction scenarios

When exploring non-random extinction scenarios, we exclusively utilized low-resolution *TR-MF_G_*curves, as the analytical formula is not applicable for them. The ASV richness range selected for the power-law regression was determined through the analysis of the random-extinction scenario. Our investigation focused on two categories of non-random extinction scenarios (Teichert et al. 2017), defining survival probability through either KO presence/absence compositional information (four *function-based* scenarios: F1-F4) or abundance distribution information (four *abundance-based* scenarios: A1-A4).

In the function-based scenarios, [F1] the *generalists survival* scenario assumed that survival probability is proportional to KO richness, reflecting a plausible situation where ASVs with greater genetic functional richness within a genome (i.e., generalists) are more resilient to environmental fluctuations (52). Conversely, [F2] the *low maintenance survival* scenario assumed that survival probability is proportional to the inverse of KO richness, attributed to the smaller maintenance cost of their genomes. [F3] The *niche uniqueness advantage* scenario assumed that ASVs with greater mean dissimilarities (evaluated by Sørensen dissimilarity measure) in KO composition compared to all other ASVs are more likely to survive. This notion is rooted in the theoretical proposition that ASVs functionally dissimilar to others may indicate less overlapping of niches, allowing them to escape competition (53). Conversely, [F4] the *shared niche survival* scenario assumed that survival probability is proportional to the inverse of the averaged dissimilarities in KO composition. While F4 may not be realized in natural environments, it was included in the scenarios to obtain pessimistic (or cautious) estimates of multifunctional redundancy. The anticipation is that scenario F4 would lead to greater impacts of ASV richness loss on *MF_G_* loss compared to other scenarios. This is because functionally unique ASVs are anticipated to go extinct first in F4, leading to a more pronounced effect on multifunctionality.

For the abundance-based scenarios, [A1] the *high mean abundance survival* scenario assumed that survival probability is proportional to the mean abundance across sites and time points, indicating a scenario where rare ASVs go extinct first (54,55). [A2] The *low abundance variation survival* scenario assumed that survival probability is proportional to the inverse of the standard deviation of abundance across sites and time points, based on the rationale that more variable ASVs are more likely to go extinct (56,57). [A3] The high *occurrence richness survival* scenario assumed that survival probability is proportional to the number of sites and time points in which the ASVs were detected, reflecting a plausible situation where ASVs widely distributed in space and time are less likely to go extinct (58–60). [A4] The *high occurrence Shannon* survival scenario assumed that survival probability is proportional to the hill number (q = 1, i.e., the 1st order effective number, Chao, Chiu and Jost (2014)) of sites and time points in which the ASVs were detected. This modification of scenario A3 considers both the occurrence and the variation in abundance.

#### Computational procedures for ASV-extinctions and multifunctional redundancy

To simulate both random and nonrandom extinction scenarios numerically, we utilized the sample() function in R to generate a shuffled sequence of ASV IDs. This sequence was created with uniform (unweighted) sampling probability for random-extinction scenario or weighted sampling probability based on survival probability for non-random extinction scenarios. In simpler terms, the final element of the generated sequence corresponded to the ASV that had gone extinct first while the first element corresponded to the one that had survived until all ASVs had gone extinct.

Following this sequence, we generated multiple ASV richness levels, covering from a single ASV to 100% of the maximum ASV richness, with increments of 5% (1 ASV, 5%, 10%, … up to 100%), resulting in 21 ASV richness levels. We repeated these procedures 100 times by preparing 100 randomly shuffled sequences as 100 trajectories of ASV extinctions. From these 100 trajectories, we calculated the mean KO richness at each ASV richness level and fitted the power-law curve using linear regression on log_10_-transformed ASV richness and mean KO richness (log10(mean KO richness) = log10(*c*) + *a*log10(ASV richness)), where the fitted exponent ’a’ served as the index of multifunctional redundancy. A smaller exponent *a* value indicated higher multifunctional redundancy.

#### Metacommunity assembly scenario

Assuming each local community within the Lake Biwa watershed originated from a shared metacommunity (36), we also employed the *TR-MF_G_* curve to investigate potential mechanisms influencing variations in multifunctionality across sites and time points. More specifically, we gathered all ASVs from local communities into a common pool as the metacommunity. Subsequently, we systematically increased ASV richness in 5% increments (ranging from 5% to 100% of total ASV richness), including one smaller value (100 ASVs), with following the survival probability defined in the extinction scenarios. Although the processes of extinction and assembly may seem to be opposites, it is essential to note that both the orders of ASV extinction and assembly can be defined by the identical survival probability. As a result, the random and nonrandom extinction scenarios can also be interpreted as the random and nonrandom assembly scenarios, respectively.

This iterative process was repeated 200 times under the random assembly scenario, generating *TR-MF_G_* curves. The 95% ranges of KO richness values at each ASV richness level were determined by extracting the 2.5% and 97.5% quantiles from the 200 replications. By overlaying the realized ASV richness and KO richness combinations, we identified local communities that deviated from the 95% range. Furthermore, we created *TR-MF_G_* curves and associated 95% ranges under non-random assembly scenarios (F1-F4 & A1-A4). We examined whether local communities showing deviations from random assembly scenarios aligned with specific non-random assembly scenarios, offering insights into the underlying mechanisms shaping the assembly processes of the focal communities.

### Computation

All computation processes that are not specified in the sections above were conducted in R (version 4.2.1) (http://www.r-project.org/). Specifically, we process sequences with the ‘dada2’ package (version 1.26.0) (47). We estimated prokaryotic diversity with the ‘iNext’ package (version 3.0.0)(62). For the linear model analysis, we used lm(), glm(), and step() functions for a simple linear regression, generalized linear model with Poisson distribution, and model selection based on Akaike Information Criterion (AIC). The R notebook as a html file especially for Ecosystem functioning experiment and ASV-extinction simulations and data files for their inputs are available at https://github.com/tksmiki/biwako_redundancy.

## Result

### Basic information of ASV richness and KO richness

After obtaining the sequencing results from the negative controls, which included 29 and 16 reads and corresponded to 12 and 7 ASVs for the extraction and PCR negative controls, respectively, we normalized the results from other data sets based on these values. Subsequently, we assessed the sampling coverages of the 20 data sets by using their sequence frequency distribution. Among them, 7 data sets exhibited sampling coverages lower than 90%, leading us to exclude them from the estimation of standardized ASV richness and KO richness. The total ASV richness and KO richness for the remaining 13 data sets are represented by 3576 and 7340, respectively, with detailed information provided in **Table S1** and **Figure S1** for each data set. One of the datasets (S11, collected from Lake Biwa on September 10, 2019) initially comprised only 2675 reads before resampling (Table S1). Although this could introduce uncertainties into subsequent analyses, its sampling coverage was sufficiently high at 96%.

Additionally, the robustness of the major statistical tests, including those associated with Figure 1 and Figure 2b, persisted even when the results from S11 were excluded.

**Figure 1:**
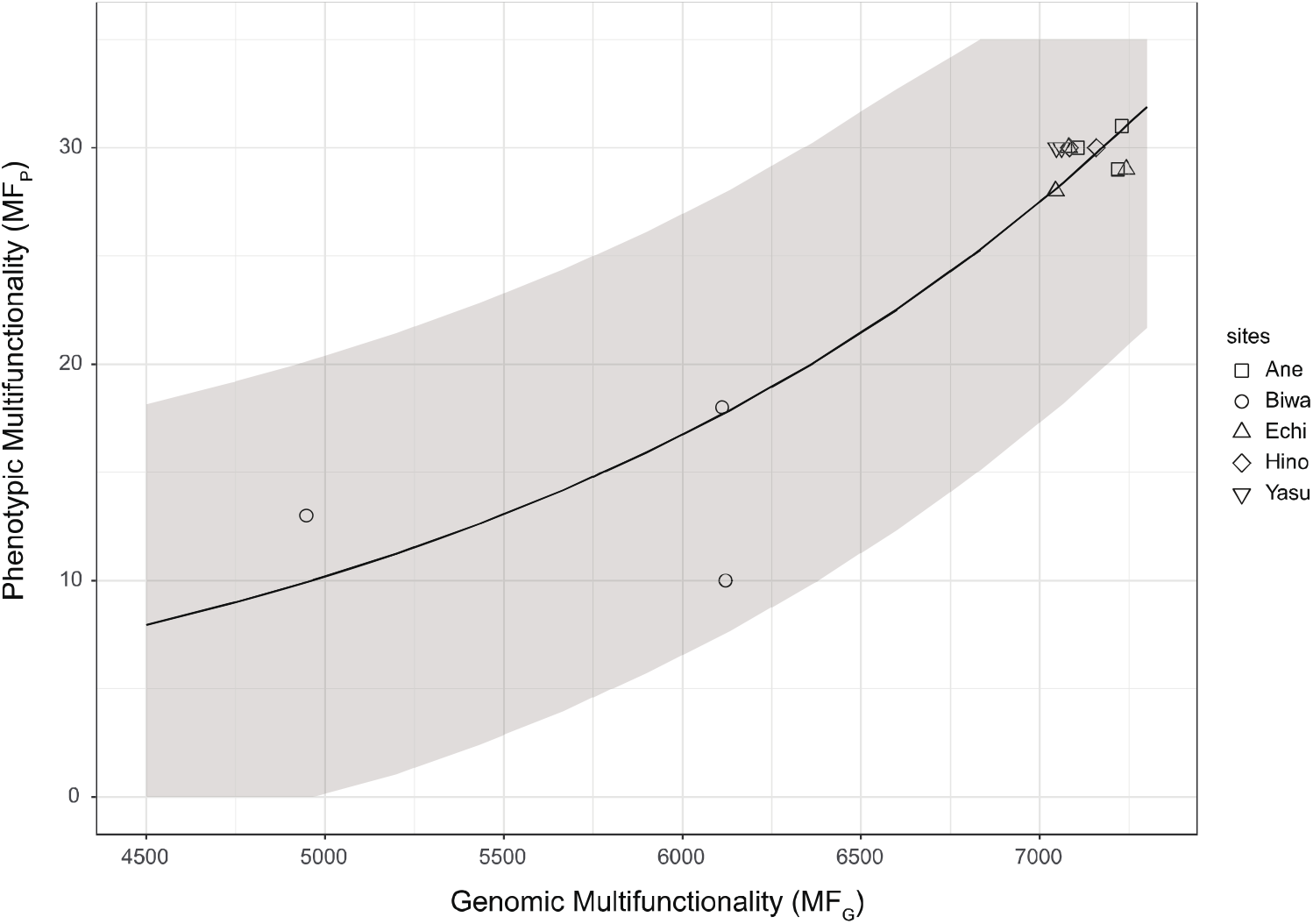
Positive association between *MF_G_* and *MF_P_*. GLM with Poisson distribution demonstrated the statistically positive association between genomic multifunctionality index (*MF_G_*) and phenotypic multifunctionality index (*MF_P_*) evaluated by Ecoplate incubation experiment. The line and shaded region represent the regression line and ±2σ ranges, respectively.

**Figure 2:**
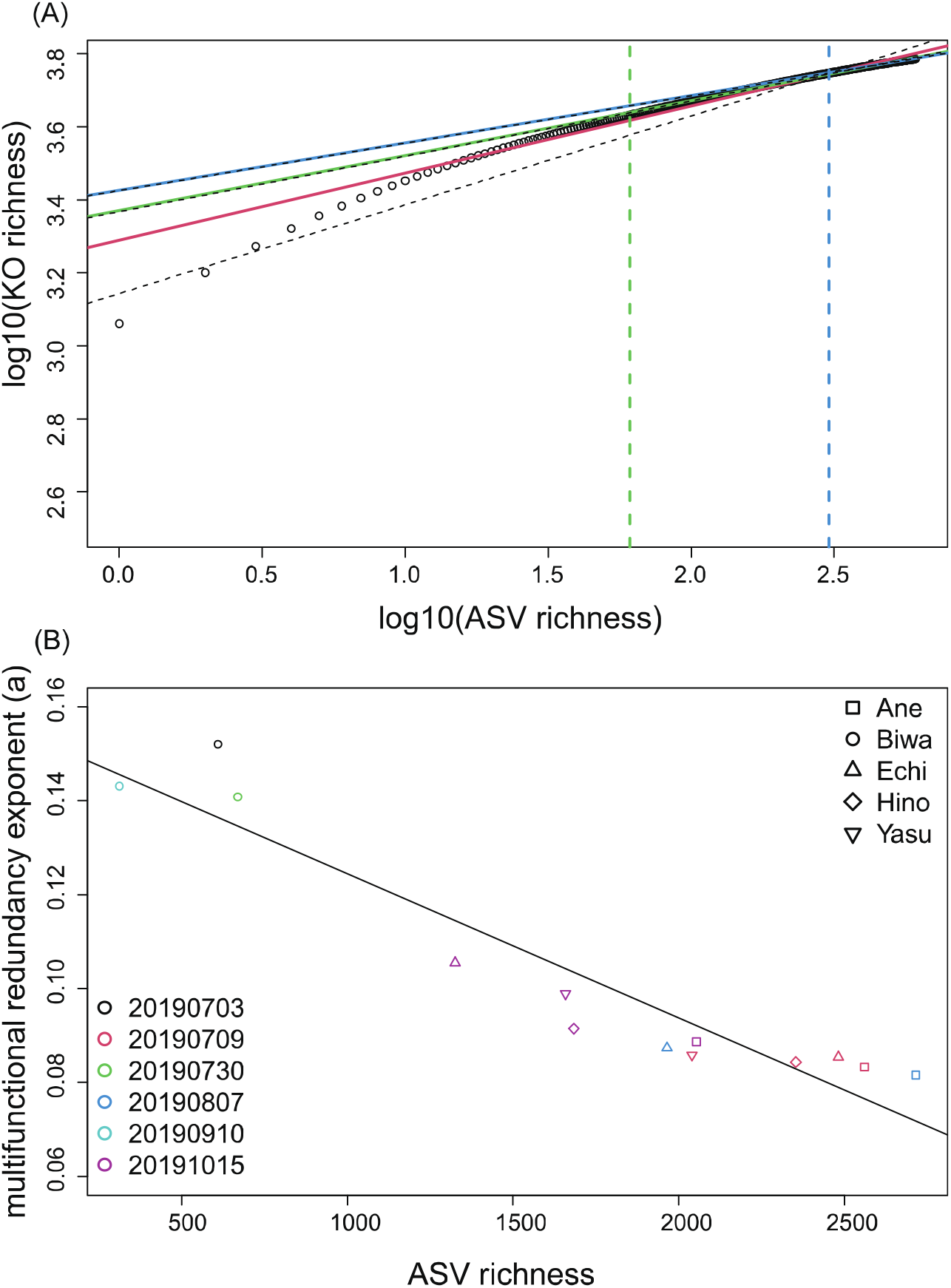
(A) Example of *TR-MF_G_* curve and its power-law regression from Lake Biwa on July 03. The points represent the analytical estimated expected KO richness for every different level of ASV richness, i.e., the high-resolution TR-MF_G_ curve. The red, light green, and light blue solids lines (or the dashed lines) represent the power-law regression on the high-resolution curve (or the low-resolution curve) from 1 ASV to 100% maximum ASV richness, 10% to 100% maximum ASV richness, and 50% to 100% maximum ASV richness, respectively. The low-resolution curve is just a subset of the high-resolution curve. Except for the fitting from 1 ASV to 100% maximum ASV richness, low-resolution regressions showed quantitatively comparable results with high-resolution regressions. The vertical light green and light blue dashed lines represent the 10% and 50 % maximum ASV richness levels, respectively. (B) Relationship between ASV richness and multifunctional redundancy exponent from the random-extinction scenario. The negative association between ASV richness and multifunctional redundancy exponent indicates the positive association between ASV richness and the magnitude of multifunctional redundancy.

### Validation and assessment of proposed methods

The generalized linear model (GLM) with a Poisson distribution revealed a significant positive linear relationship between *MF_G_* and *MF_P_* (λ = 0.0004963*MF_G_ – 0.1611650, P value for the coefficient of *MF_G_* = 1.67e-5), indicating a positive association between the two variables (**Fig. 1**). Utilizing the step() function to identify the best model explaining the variation in *MF_P_* with potential explanatory variables (ASV richness and *MF_G_*), we observed that only *MF_G_* remained as a significant explanatory variable. This indicated that genomic multifunctionality (*MF_G_*) is a good proxy of phenotypic multifunctionality (*MF_P_*).

The power-law fitting applied to high- and low-resolution *TR-MF_G_*curves revealed that the estimated intercepts (*c*) for the entire range of ASV levels (from a single ASV to the maximum ASV) were considerably larger than the expected MF_G_ with a single ASV (**Fig. 2a** and **Table S2**), indicating a tendency for multifunctionality overestimation when TR is very low. Analyzing the estimates of the exponent (*a*) across three defined ranges, we found that two intervals, specifically from 10% to 100% and from 50% to 100% of the maximum ASV richness, yielded quantitatively comparable results for *a* between the high- and low-resolution curves. Considering these findings, we opted to utilize the range from 10% to 100% of the maximum ASV richness for numerical simulations under non-random extinction scenarios, ensuring coverage of a broader interval, provided that the estimates from the low-resolution curves remain comparable to those from the high-resolution curves (within 3% differences, **Table S2**).

Utilizing estimates obtained from the selected ASV richness ranges (10% - 100% of the maximum ASV richness for each community) under the random extinction scenario, we identified variations in multifunctional redundancy across different sites and sampling dates. Notably, we observed a positive association between these variations and ASV richness (**Fig. 2b**), where a smaller exponent value indicated greater multifunctional redundancy (linear regression, adjusted R^2^ = 0.8838).

### Variations in multifunctional redundancy under non-random extinction scenarios

The redundancy exponent (a) exhibited substantial variations across non-random extinction scenarios (**Fig. 3, Fig. 4**), although the ranking order of the exponent among sites and dates remained generally consistent between scenarios (**Fig. 4a**). In comparison to redundancy estimated from random-extinction scenarios, F1 and F3 demonstrated higher redundancy (i.e., smaller exponent), while F2, F4, and A1 exhibited lower redundancy (i.e., greater exponent)(**Fig. 4b**). Notably, scenario F2, representing the *low maintenance survival* scenario, estimated the lowest redundancy, reflected by the highest exponent. The other scenarios did not exhibit a clear trend in redundancy estimation.

**Figure 3:**
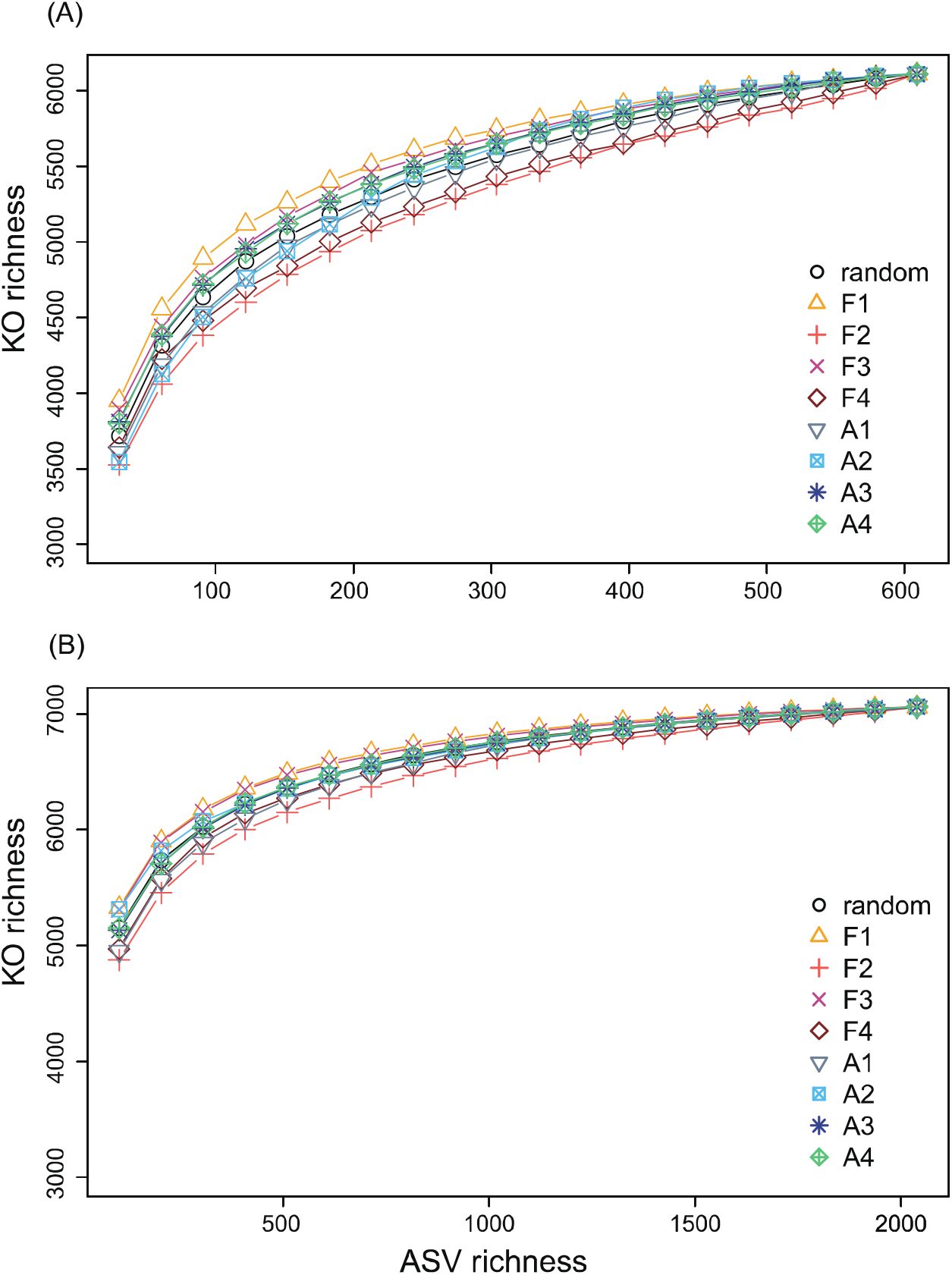
Examples of TR-MF_G_ curves from all scenarios. (A) The dependence of KO richness reduction along with different ASV extinction scenarios for the community collected on July 03 from Lake Biwa. (B) The dependence of KO richness reduction along with different ASV extinction scenarios for the community collected on July 09 from Yasu river.

**Figure 4:**
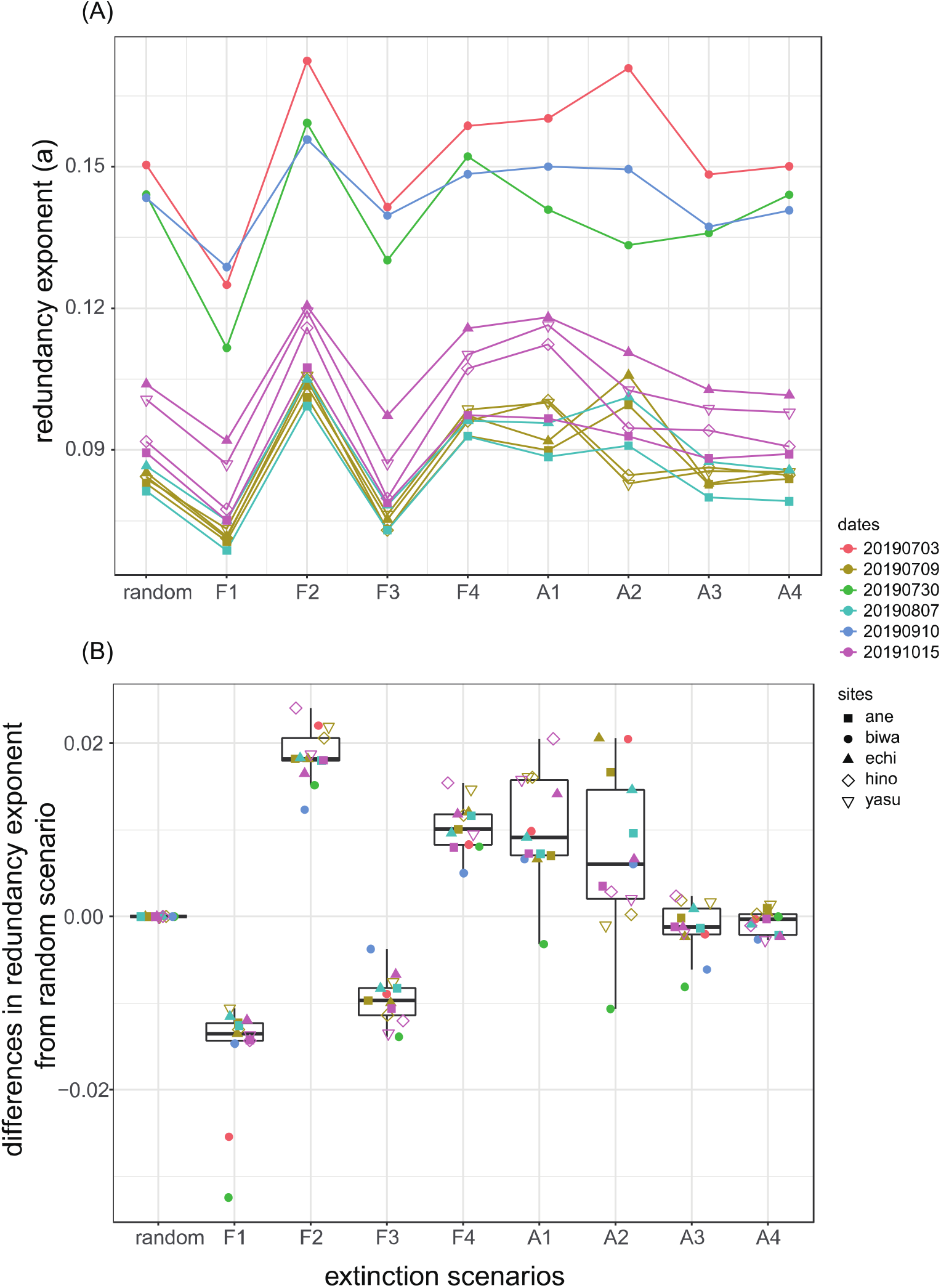
Variations in redundancy exponents between scenarios. Summary of the dependency of the redundancy exponent (*a*) on extinction scenarios, sites, and time points. (a) The direct comparison of the redundancy exponent; the greater exponent value represents lower multifunctional redundancy. (b) The differences between the exponent values between those from the random extinction scenario and those from eight non-random extinction scenarios. The positive (or negative) values represent the lower (or higher) multifunctional redundancy under the focal non-random extinction scenario than that under the random-extinction scenario.

### Comparison of multifunctionality through community assembly from a shared metacommunity

We observed that the majority of communities within the metacommunity can be explained by the random assembly scenario, as indicated by the 95% range. Nevertheless, four local communities (Biwa on 0703, 0730, and 0910, and Echi on 1015) deviated from this95% range of random assembly (**Fig. 5**). One of them (Echi on 1015) could be explained by F1 and F3. Additionally, the deviations of two communities (Biwa on 0703 and 0730,) were explainable by scenario F2, with one of them (Biwa on0703) also aligning with scenario F4 (but at the boundary of the 95% range). One local community (Yasu on 0709) also fell into A2 (**Fig. S2**). One local community (Biwa on 0910) did not fall into any scenarios (random, F1-F4, and A1-A4).

**Figure 5:**
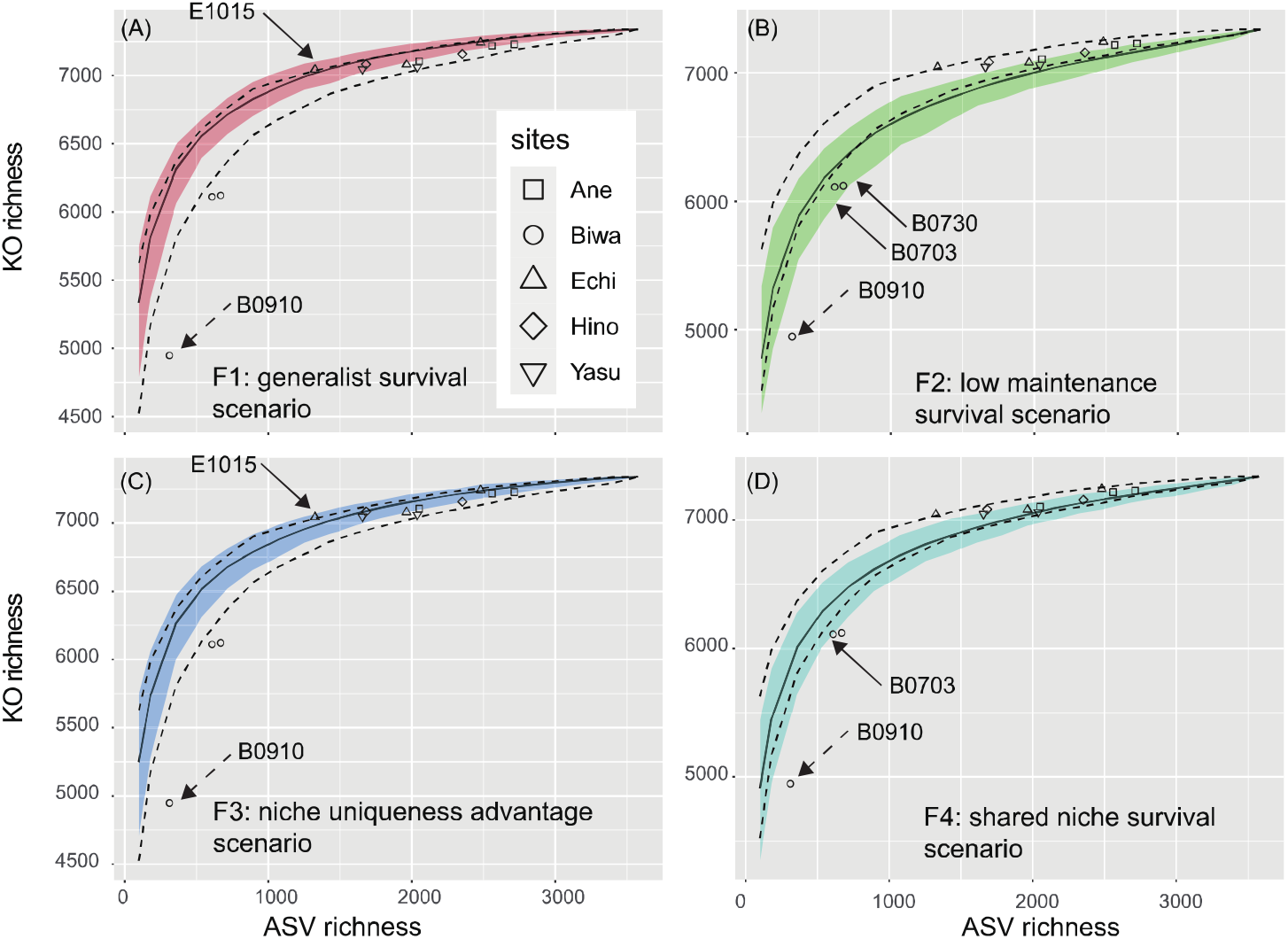
Deviation from random-assembly from the metacommunity for function-based scenarios (F1-F4). The variations in ASV richness and KO richness between 13 communities and their relationship with the random-assembly (i.e. random-extinction) scenarios’ 95 % confidence interval from 200 repeated simulations (the range between the dashed lines) and with the 95% confidence interval (color-shaded region) of one of the four function-based nonrandom-extinction scenarios (F1-F4). Communities indicated by the solid arrows (i.e., B0703, B0730, and E1015) correspond to the communities from Lake Biwa on July 3, July 30, and from Echi river on October 15, respectively. These communities cannot be explained by random assembly processes but can be attributed to a part of the non-random assembly scenarios (F1-F4). Community indicated by dash arrow (i.e., B0910, Community from Lake Biwa on September 10) cannot be explained by any scenarios (random, F1-F4, and A1-A4).

## Discussion

### Advancements in Methodology for Assessing Multifunctional Redundancy

The primary objective of our study was to refine the methodology for assessing multifunctional redundancy, resulting in three key findings with significant implications.

Firstly, the implementation of PICRUSt2 was indirectly validated through a comparison of KO richness predictions with phenotypic multifunctionality assessed by ecoplate incubations (**Fig. 1**). Unlike the previous study (21), our analysis covered a wide bacterial richness range (from 312 to 2715 ASVs), demonstrating the practical effectiveness of our proposed procedure starting from amplicon sequences. Model selection further revealed that KO richness serves as a superior predictor of phenotypic multifunctionality compared to ASV richness. Although direct metagenomic sequencing for validation is acknowledged, these results highlight the practical efficacy of our proposed approach for obtaining functional information from amplicon sequences.

Secondly, our detailed analysis of the power-law regression method led to two crucial procedural enhancements. Firstly, we recommended setting the focal range for power-law regression as 10% to 100% of the maximum ASV richness to avoid dependence on the resolution of ASV richness. This adjustment ensures that the estimate is robust across different resolutions. Additionally, we proposed conducting low-resolution simulations (e.g., every 5% of the maximum ASV richness) due to computational efficiency, while confirming that power-law fittings on low-resolution *TR-MF_G_*curves are quantitatively equivalent to those on high-resolution curves. The power-law regression tends to overestimate *MF_G_* when ASV richness is low, limiting its utility for predicting the impacts of severe richness reduction on ecosystem multifunctionality. When predicting such impacts, we recommend directly using the results from extinction simulations instead of relying on power-law fitting estimates.

Lastly, concerning extinction scenarios, we proposed that random-extinction, generalists survival, and low maintenance survival scenarios represent the minimum requirements to cover the possible range of functional redundancy estimates (**Fig. 4**). The random-extinction scenario acts as the baseline, while the generalists survival and low maintenance survival scenarios provide the highest and lowest redundancy estimates (smallest and greatest exponent values), respectively, among all scenarios. This comprehensive approach ensures a thorough understanding of the potential range of functional redundancy estimates under different extinction scenarios.

### Comparative Analysis of Lake Biwa and Inlet Rivers: Multifunctional Redundancy Dynamics

The secondary objective of our study was to apply the refined methodology to multiple systems in the Lake Biwa watershed, elucidating spatio-temporal variability in multifunctional redundancy.

Three major findings emerge from our analysis: Firstly, Lake Biwa exhibited smaller bacterial richness compared to any of the four inlet rivers, irrespective of the time points (**Fig. 2**). One might argue that the observed lower richness in Lake Biwa could be attributed to a smaller number of sequences or lower bacterial abundance. However, the former is not supported as the sampling coverages were high enough (> 96%, as shown in **Table S1**). The latter is also not supported because ASV richness in rivers with comparable bacterial abundance was much greater, and temporal variations in ASV richness were much smaller than those of bacterial abundance (**Fig. S3**).

Instead, this result aligns with the established notion that high richness is likely linked to elevated environmental and spatial heterogeneity, providing multiple niches for diverse microbes to inhabit in rivers (63). Although environmental variables were not directly measured, this hypothesis gains indirect support from the relationship between *MF_G_* and *MF_P_* (Fig 1), indicating higher functional diversity in rivers compared to lakes.

Secondly, we observed a negative association between bacterial richness and the exponent of multifunctional redundancy, suggesting reduced multifunctional redundancy with a decline in bacterial richness. (**Fig. 2b**). As Lake Biwa exhibited smaller bacterial richness and redundancy compared to any of its inlet rivers, irrespective of the time points (**Fig. 2b**), it highlights a higher conservation priority for Lake Biwa compared to the four inlet rivers, at least concerning prokaryote-mediated ecosystem multifunctionality. Conversely, while it appeared that the Echi River had the lowest redundancy (greatest exponent) on July 9 and October 15, all four rivers displayed significant fluctuations in both bacterial richness and multifunctional redundancy across different time points. This complexity makes it challenging to definitively identify the river most vulnerable to disturbances and losses in bacterial taxonomic richness, affecting ecosystem multifunctionality.

The challenge in prioritizing ecosystem vulnerability between four rivers (Ane, Echi, Hino, and Yasu) may stem from the non-intensive sampling design (only four time points from each river). Additionally, the lack of clear differences between these rivers could be influenced by random assembly processes, suggested by the metacommunity scenario (**Fig. 5**). Most bacterial communities in these rivers seemed to follow a random-assembly scenario. If true, significant differences in terms of multifunctional redundancy between these four rivers may not exist. To confirm the presence or absence of real differences, more frequent sampling with additional time points and multiple locations within a river is necessary (64). Additionally, expanding the study to encompass a broader selection of inlet rivers is crucial, given the presence of 117 first-class inlet rivers in the Lake Biwa watershed.

Finally, the results suggest that Lake Biwa was subject to different assembly processes compared to its inlet rivers (**Fig. 5**). While most of the rivers could be explained by random or generalist survival scenarios, Lake Biwa aligns more closely with a low maintenance survival scenario, indicating that generalists are less likely to persist. Such contrasting results may be attributed to the distinct environments of rivers and lakes (65). For example, the running water in the inlet rivers is highly dynamic and fluctuating, which would favor generalists more because they can adapt to various conditions (66–70). In contrast, pelagic water in a large lake exhibits characteristics of a more stable environment. Therefore, the high maintenance cost for multiple genes might be a challenge for species to survive in lakes, resulting in the survival of specialists with low maintenance costs.

### Advancing methodology: underexplored aspects and future directions

In this section, we highlight three underexplored aspects of our proposed method, each serving as potential avenues for future research.

Firstly, our initial expectation that the scenario F4 would lead to lower multifunctional redundancy compared to the scenario F2 was contradicted by the results (**Fig. 4**). While F4 assumes that ASVs with greater functional dissimilarity go extinct earlier, resulting in a greater negative impact on community-wide KO richness than F2, the opposite trend was observed. A simple linear association between the indices for F2 (inverse of KO richness) and F4 (inverse of mean dissimilarity) was not evident (**Fig. S4a**). One potential refinement could involve a more direct method for defining functional uniqueness, such as considering the number of unique KOs not shared with other ASVs. However, our exploration revealed only 127 ASVs with such unique KOs, and the maximum number of unique KOs per ASV was limited to 16 (not shown). This indicates that such an index may not be effective in weighting extinction probability across 3576 ASVs. In the scenario F2, we counted the number of different KOs without considering multiple copies, but an alternative approach could involve considering these copies as well. Future investigations should also explore alternative extinction scenarios capable of yielding lower multifunctional redundancy (greater exponent *a*) than those from F2 and F4. Conservative estimates of redundancy are crucial for assessing ecosystem vulnerability, particularly when faced with substantial uncertainty. This challenge is inherent because we primarily rely on genomic information, including PICRUSt2 predictions (as in this study) or metagenome-assembled genomes (71,72), as well as abundance distribution information, but have limited access to direct physiological information of unculturable bacteria from environmental samples. Further refinement of extinction scenarios and indices is warranted to enhance the accuracy and reliability of redundancy assessments in the face of such complexities.

Second, the comparison of redundancy exponents between this study and past studies (21,29) provides insights into the robustness of the proposed method and offers cautions for its application to other systems. The exponent (a) values estimated in Miki, Yokokawa and Matsui (2014) ranged from 0.55 to 0.75, but these may have been overestimated as they were derived from pseudo-communities containing a single strain from each genus. In natural environments, where multiple strains, OTUs, or ASVs from each genus coexist in local communities, functional overlap within communities is likely higher. When applying a power-law regression directly fitting on multiple datasets, our results yielded an exponent (*a*) of 0.15799 (adjusted R^2^ = 0.9022) (**Fig. S5**), comparable to the value (*a* = 0.1338) presented in Fig.3d of Ruhl et al. 2022. However, it is important to note that our study used KO richness as a proxy for genomic multifunctionality, while Ruhl et al. used Pfam richness. Additionally, the unit of taxonomic richness (OTUs or ASVs) also plays a crucial role, as highlighted by Ruhl et al. 2022, indicating that the estimated exponent highly depends on both the proxy of genomic multifunctionality and the taxonomic richness unit (Fig. 3 of Ruhl *et al.* 2022).

The third aspect, while not directly aligned with the main objectives of this study, offers the potential for a more in-depth analysis of the relationship between the predicted KO composition, its richness, and the spatio-temporal distribution of each ASV in the watershed. In line with the exploration of utilizing genomic functional information for predicting bacterial occurrence patterns, as proposed by studies like Barberán *et al.* (2014), we also found a positive yet weak association between KO richness (indicative of functional generalization) and both mean abundance (**Fig. S4b**, linear regression on log10-transformed mean abundance, slope = 4.589e-04 (P < 2.0e-16), adjusted R^2^ = 0.04647) and the number of occurrences (**Fig. S4c**, linear regression, slope = 0.0020760 (P < 2.0e-16), adjusted R^2^ = 0.0632). While KO richness of ASVs is not directly linked to their genome size, these patterns stand in contrast to observations in the surface ocean and lakes, where genome-streamlined groups tend to dominate (74–76). To maintain the simplicity of the multifunctional redundancy assessment procedure, we exclusively utilized KO richness without incorporating specific functional information for each KO. However, introducing functional details of KO composition into the procedure could yield two significant advantages: 1) refining the potential scenarios of bacterial richness loss, thereby reducing uncertainty in the estimated redundancy exponent, and 2) providing more detailed assessments of functional redundancy, such as redundancy within specific functional categories like carbon metabolism and nitrogen metabolism. In the context of this study’s primary objectives, maintaining simplicity in the assessment of multifunctional redundancy was prioritized. However, for researchers willing to employ more complex procedures, exploring predictions of growth rate and interspecific interactions through metabolic network-based reverse ecology methods, such as flux balance analysis (FBA), can be a promising avenue. It’s worth acknowledging that these tools are still evolving in their development (77,78).

In light of these underexplored features and the current limitations in the size of amplicon sequence data, future investigations may prioritize refining our methodology, addressing potential complexities, and collecting spatially and temporally high-resolution datasets. The crucial refinement of our methodology is necessary for accurately assessing the vulnerability of ecosystem functions, particularly those mediated by microorganisms, and will contribute to the development of effective strategies for ecosystem management and conservation.

## Acknowledgements

This study was supported by Center for Ecological Research, Kyoto University, a Joint Usage / Research Center. We appreciate Captain Dr. Y. Goda and Vice-Captain T. Akatsuka of the Research vessel Hasu for field sampling in Lake Biwa, and R. Nakamura and A. Matsuda for helping field samplings in the rivers.

## Funding

T.M., H.Y., and S.N. were supported by Grant for Environmental Research Projects, Sumitomo Foundation. T.M. and T.Y were supported by JSPS KAKENHI, Grant-in-Aid for Scientific Research (S) (19H05667). T.M. and K.Y were supported by JSPS KAKENHI, Grant-in-Aid for Scientific Research (A) (23H00538). T.M and N.S. were supported by JSPS KAKENHI, Grant-in-Aid for Scientific Research (B) (19H03302). T.M. was also supported by JSPS KAKENHI, Grant-in-Aid for Scientific Research (A) (19H00956) and the Alexander von Humboldt fellowship.

**Authors’ contributions**

WHC and TM leads data analysis, design and write the manuscript, MI leadingly conducted sampling and experiments, KY, KM, and HY helped experimental procedures. TM, KM, TY, HY, and SN developed the whole picture of the project in the lake Biwa watershed and contribute to the improvement of the manuscript.

