## Supplementary Data for "Advancing marker-gene-based methods for prokaryote-mediated multifunctional redundancy: exploring random and nonrandom extinctions in a watershed"

Title:

**Supplementary methods**

DNA extraction

In the initial phase of DNA extraction, designed to prevent mechanical damage to the filter cartridge and minimize contamination risks, we employed vacuum (EZ-Zac Vacuum Manifold) to remove the RNAlater solution from each filter cartridge, followed by a gentle wash with 1 mL of MilliQ water. Subsequent DNA extraction employed the DNeasy® Blood & Tissue Kit (QIAGEN, Hilden, Germany), following the protocol outlined by Miya et al. (2016). In brief, a mixture of proteinase K solution (20 μL), PBS (220 μL), and buffer AL (200 μL) was prepared, and 440 μL of this mixture was added to each filter cartridge. Lysis of materials on the cartridge filters occurred during a 10-minute incubation on a rotary shaker (15 rpm) at 56 °C. Following incubation, filter cartridges underwent vigorous shaking with zirconia beads inside for 3 minutes (3,200 rpm; Vortex-Genie 2, Scientific Industries) (Ushio 2019). The resulting mixture was then transferred into a new 2 ml tube from the filter cartridge inlet through centrifugation (3,500 × g for 1 minute). Purification of the collected DNA employed the DNeasy® Blood & Tissue Kit according to the manufacturer's protocol. Post-purification, the DNA was eluted using 100 μL of the provided elution buffer. Throughout these procedures, an extraction negative control was prepared to monitor potential contamination. The eluted DNAs were stored at –20 °C until further processing.

PCR amplification and sequencing

Before starting library preparation, we sterilized workspaces and equipment. Filtered pipette tips were used, and we separated pre- and post-PCR samples to prevent cross-contamination. A single negative control (PCR-negative control) was included to monitor contamination throughout PCR procedures (the sample S021 and S022; see [3.1.4] in sumitomo2024_main.nb_20240222.html at <https://github.com/tksmiki/biwako_redundancy>). In the first PCR step, we combined 6 μL of 2 × KAPA HiFi HotStart ReadyMix (KAPA Biosystems, Wilmington, WA, USA), 0.7 μL of prokaryotic universal primer (eurofins, 5 μM for each primer; Forward, 515F (5’-[illumina MiSeq adapter]-NNNNNN-GTGYCAGCMGCCGCGGTAA-3’ (Parada et al. 2016); Reverse, 806R (5’-[illumina MiSeq adapter]-NNNNNN-GGACTACNVGGGTWTCTAAT-3’) (Apprill et al. 2015)), 2.6 μL of MilliQ water, and 2 μL of DNA template. The thermal cycling profile began with an initial 3-minute denaturation at 95 °C, followed by 35 cycles of denaturation at 98 °C for 20 seconds, annealing at 60 °C for 15 seconds, and extension at 72 °C for 30 seconds. A final extension step was performed at 72 °C for 5 minutes. For purification, the first PCR product (18 μL each) underwent purification using AMPure XP beads (PCR product: AMPure XP beads = 1:0.8; Beckman Coulter, Brea, California, USA). The resulting purified products were then 10-fold diluted and utilized as templates for the second PCR.

The second PCR was conducted with a 24 μl reaction volume, comprising 12 μl of 2 × KAPA HiFi HotStart ReadyMix, 1.4 μl of each primer (5 μM of each primer), 7.2 μl of sterilized distilled H_2_O, and 2.0 μl of template. Various combinations of forward and reverse indices were employed for distinct templates (samples) to facilitate massive parallel sequencing with MiSeq. The thermal cycle profile initiated with an initial 3-minute denaturation at 95 °C, followed by 12 cycles: denaturation at 98 °C for 20 seconds, annealing at 68 °C for 15 seconds, and extension at 72 °C for 15 seconds, concluding with a final extension at 72 °C for 5 minutes.

Each 20 μL second PCR product was combined into a pooled library. This library was subsequently purified using AMPure XP beads (PCR product: AMPure XP beads, 1:0.8 ratio). Size selection involved isolating fragments peaking between 400-500 bp using 2% E-Gel Size Select (Thermo Fisher Scientific, Waltham, MA, USA). The concentration of double-stranded DNA in the library was quantified using a Qubit dsDNA HS assay kit and Qubit fluorometer (Thermo Fisher Scientific, Waltham, MA, USA). The concentration was adjusted to 4 nM using Milli-Q water, and the DNA was prepared for sequencing on the MiSeq platform (Illumina, San Diego, CA, USA). Sequencing was carried out using a MiSeq Reagent Nano Kit v2 for 2 × 250 bp PE (Illumina, San Diego, CA, USA). Details on the sequence depth of each sample are provided in Table S1.

Computation

Major computation processes were summarized as the R Notebook html file at <https://github.com/tksmiki/biwako_redundancy>.

(Continued to the next page)

**Supplementary Figures**

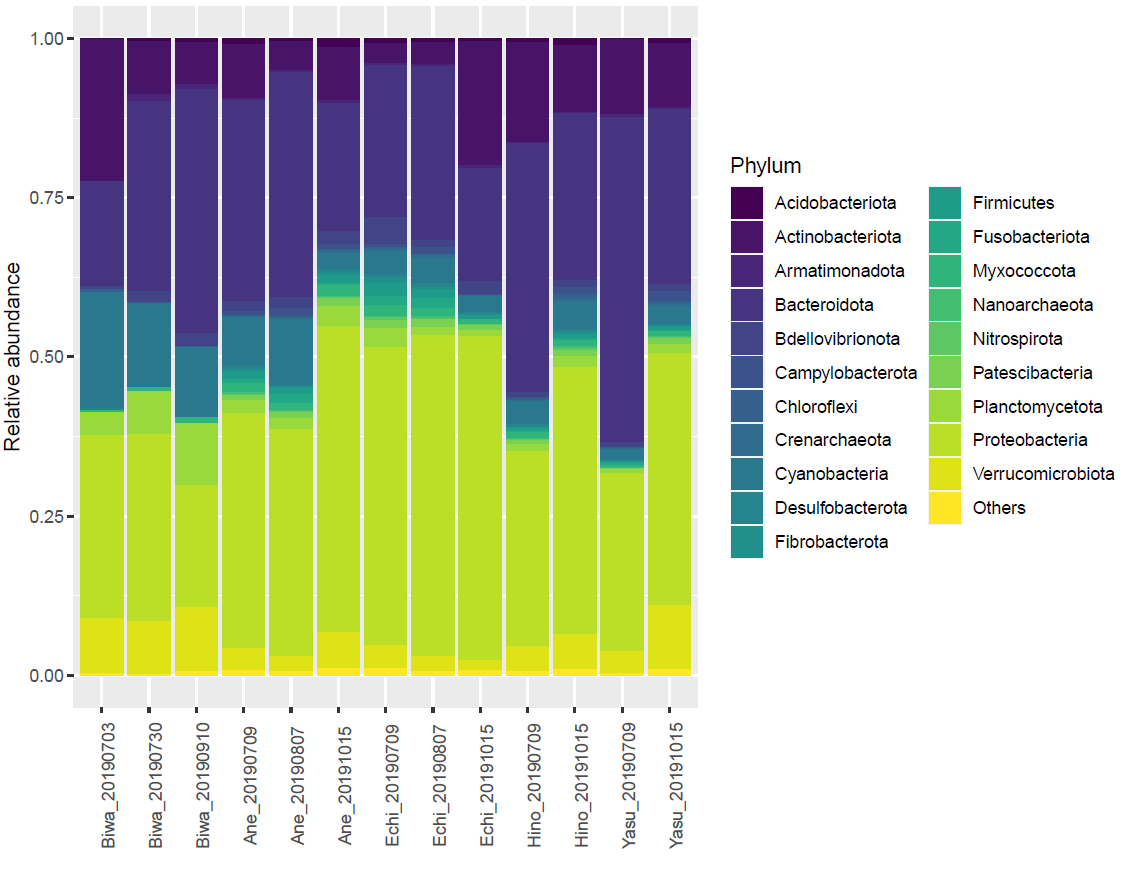

**Figure S1: The taxonomic composition for each sampling site.** The mean abundance out of the top 20 abundant phyla are grouped and labeled as "Others".

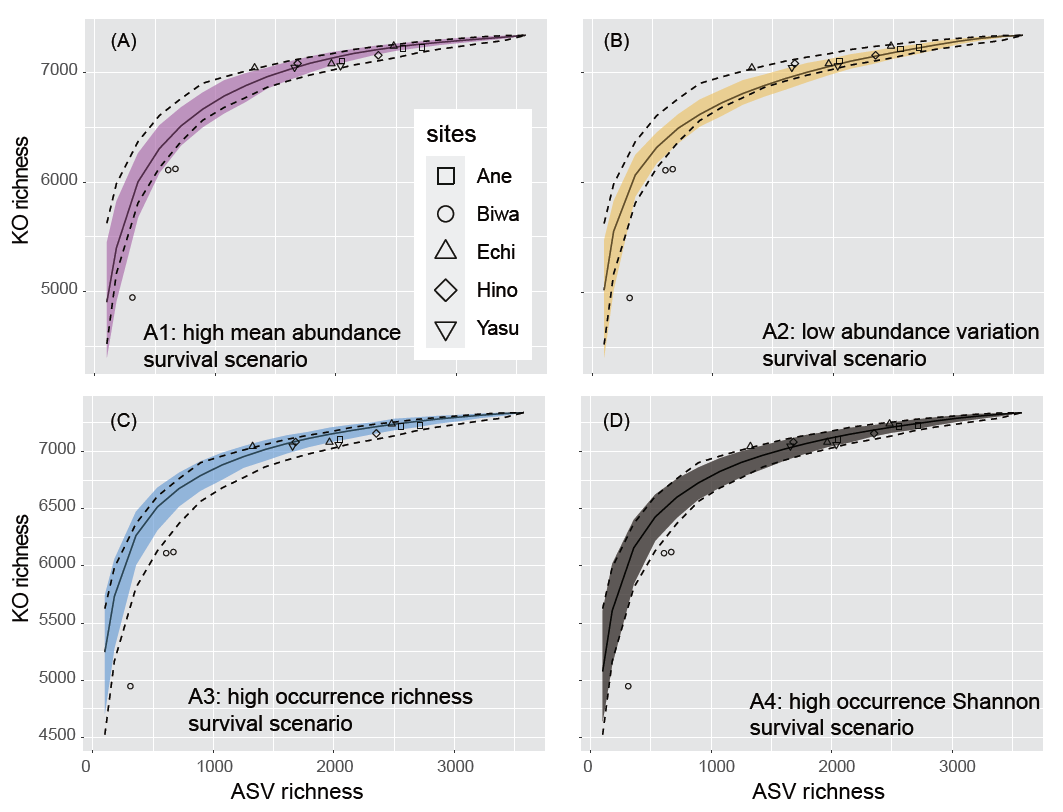

**Figure S2: Deviation from random-assembly from the metacommunity for abundance-based scenarios (A1-A4).** The variations in ASV richness and KO richness between 13 communities and their relationship with the random-assembly (i.e. random-extinction) scenarios’ 95 % confidence interval from 200 repeated simulations (region between the dashed lines) and with the 95% confidence interval (color-shaded region) of one of the four abundance-based nonrandom-extinction scenarios (A1-A4). Any of four communities (B0703, B0730, B0910, E1015) did not fall into these scenarios (A1-A4).

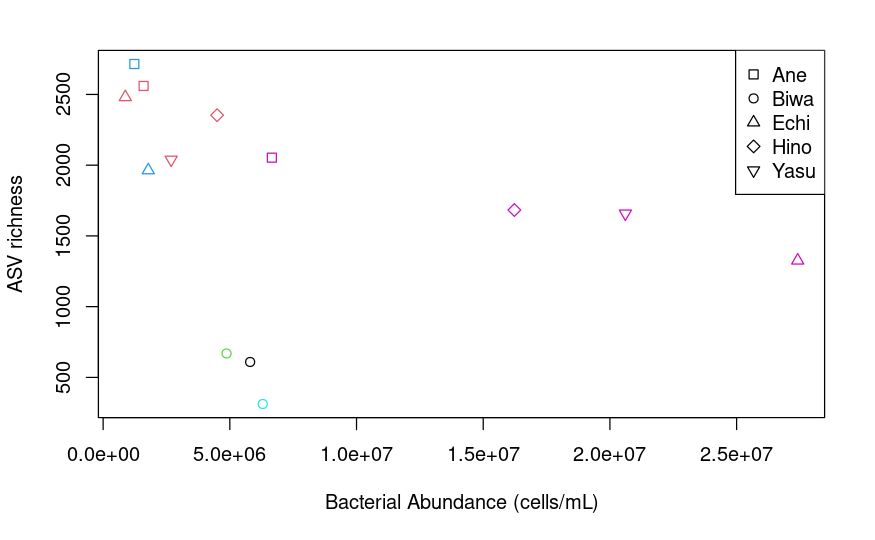

**Figure S3:** Bacterial abundance and ASV richness. The color setting is identical to Fig. 2b.

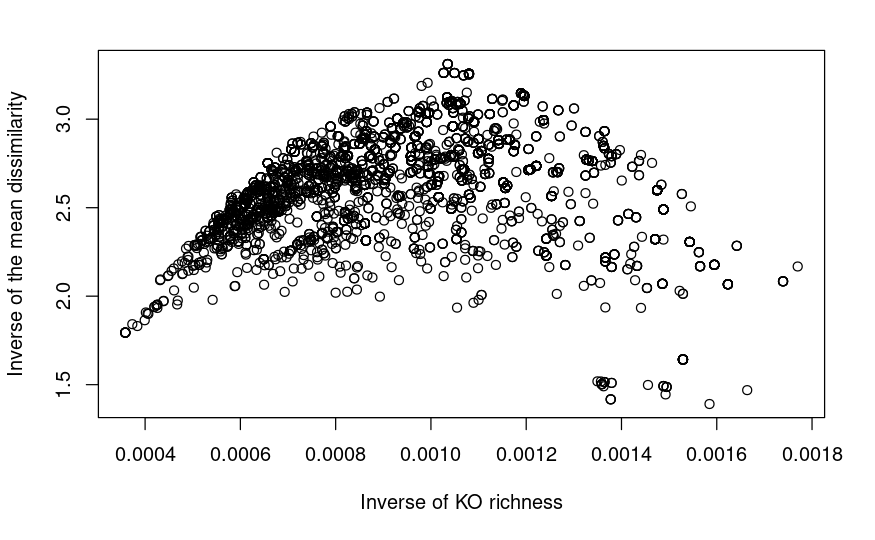

**Figure S4a: Nonlinear association between the genomic cost index and functionally uniqueness index.** The horizontal and vertical axes represent the relative survival probability of ASVs based on genomic cost scenario (i.e., proportional to the inverse of KO richness) and that of ASVs based on functionally uniqueness scenario (i.e., proportional to the inverse of the mean dissimilarity of KO composition).

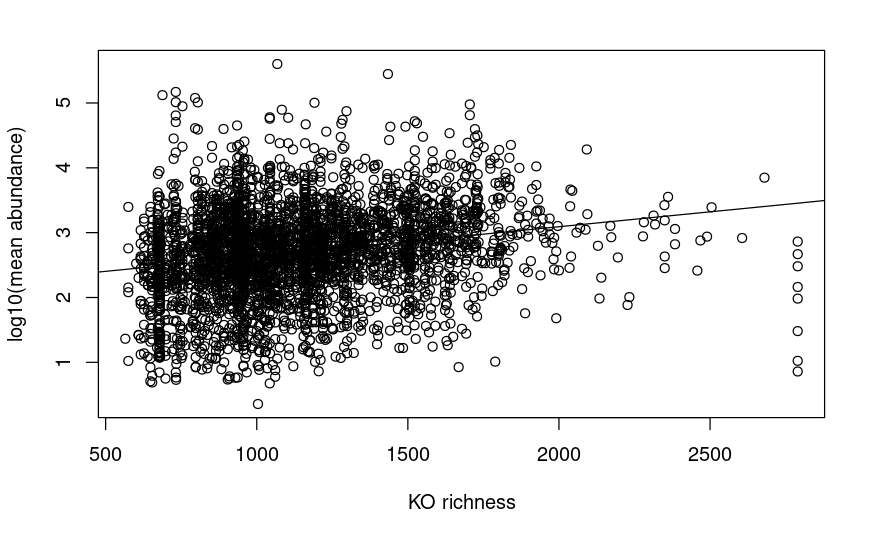

**Figure S4b: Positive association between the functional generality index and log-transformed mean abundance.** The statistically significant but weak positive association between two variables (KO richness and 1og-10 transformed mean abundance).

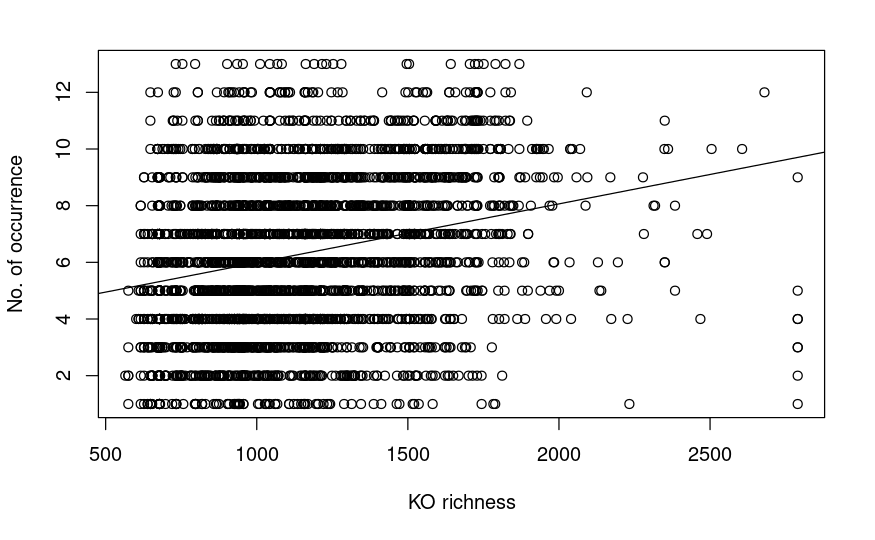

**Figure S4c: Positive association between the functional generality index and the number of occurrence (1-13).** The statistically significant but weak positive association between two variables (KO richness and the number of occurrence).

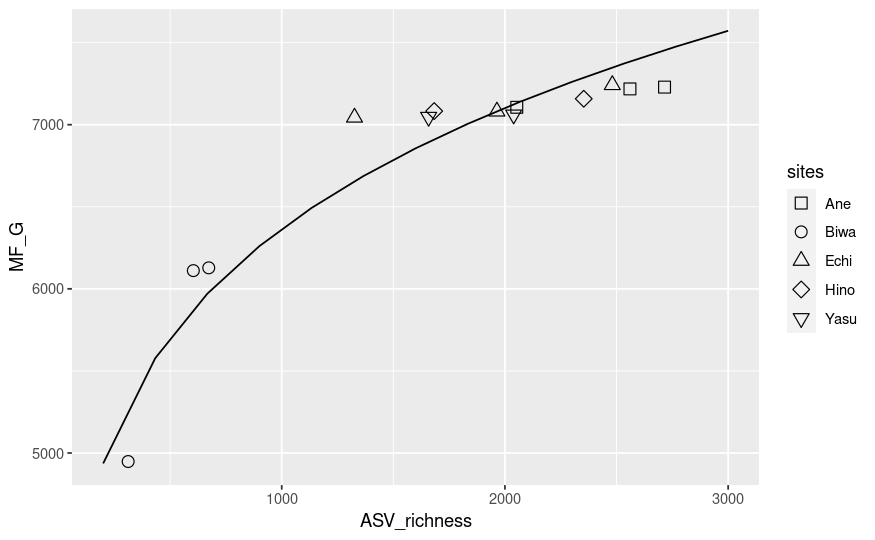

**Figure S5: Direct fitting of power-law regression on all data sets (all local communities) from the Lake Biwa watershed.** The fitted function the power-law regression, i.e., log10 transformed linear regression is (*a* = 0.15779, *c* = 10^3.32986^ = 2137.27, adjusted R^2^ = 0.9022).

**Supplementary Tables**

In the excel file (TablesS1_S2.xlsx):

**Table S1: The list of #reads, observed ASV richness, coverage for all samples, & standardized ASV richness, estimated KO richness for the remaining data sets.**

**Table S2: List of statistics (point and interval estimates of c & a, the expected KO richness at a single ASV richness, and adjusted-R squared)**
